# High density linkage to physical mapping in a unique Tall x Dwarf Coconut (*Cocos nucifera* L.) outbred F_2_ uncovers a major QTL for flowering time colocalized with the *FLOWERING LOCUS T (FT)*

**DOI:** 10.1101/2024.01.31.578260

**Authors:** Dario Grattapaglia, Wellington Bruno dos Santos Alves, Cleso Antônio Patto Pacheco

## Abstract

**Introduction:** The coconut tree crop (*Cocos nucifera* L.) provides vital resources for millions of people worldwide. Coconut germplasm is largely classified into ‘Tall’ (Typica) and ‘Dwarf’ (Nana) types. While Tall coconuts are outcrossing, more productive, stress tolerant and late flowering, Dwarf coconut are inbred, flower early with a high rate of bunch emission. Precocity determines earlier production of a plantation and facilitates management and harvest.

**Methods:** We used a unique outbred F_2_ population generated by intercrossing F_1_ hybrids between Brazilian Green Dwarf from Jiqui (BGDJ) and West African Tall (WAT) cultivars. SNP markers fixed for alternative alleles in the two varieties, segregating in an F_2_ configuration were used to build a high-density linkage map with ∼3,000 SNPs, anchored to the existing chromosome-level genome assemblies, and a QTL mapping analysis was carried out.

**Results:** The linkage map established the chromosomes numbering correspondence between the two reference genome versions and the relationship between recombination rate, physical distance and gene density in the coconut genomes. Leveraging the strong segregation for precocity inherited from the Dwarf cultivar in the F_2_, a major effect QTL with incomplete dominance was mapped for flowering time. *FLOWERING LOCUS T* (*FT*) gene homologs of coconut previously described as putatively involved in flowering time by alternative splice variant analysis, were colocalized within a ∼200kb window of the major effect QTL (LOD=11.86).

**Discussion:** Our work provides strong phenotype-based evidence for the role of the FT locus as the putative underlying functional variant for the flowering time difference between Dwarf and Tall coconuts. Major effect QTLs were also detected for developmental traits of the palm, plausibly suggesting pleiotropism of the *FT* locus for other precocity traits. Haplotypes of the two SNPs flanking the flowering time QTL inherited from the Dwarf parent BGDJ caused a reduction in the time to flower of around 400 days. These SNPs could be used for high throughput marker assisted selection of early flowering and higher productivity recombinant lines, providing innovative genetic material to the coconut industry.

## 1. Introduction

The coconut palm (*Cocos nucifera* L.) is a worldwide important tropical tree crop providing vital resources for millions of farmers and industries with a diverse use of its oil and fruit products. The only species belonging to the genus *Cocos*, the coconut palm is a diploid (2n = 32) monoecious, perennial monocotyledon of the family *Arecaceae* (Palmae), with a relatively large genome size (1C = 2.74 Gb). With its most likely center of origin in Southeastern Asia, it is currently cultivated in more than 92 tropical countries (Batugal et al., 2009; Gunn et al., 2011) covering more than 12 million hectares worldwide (FAO, 2021).

Coconut germplasm is largely classified into ‘Tall’ (Typica) and ‘Dwarf’ (Nana) types. Tall, a.k.a. Giant coconuts, are preferentially outcrossing, genetically diverse and display high productivity, broad adaptability and tolerance to specific pests and diseases. Dwarf coconuts are mostly self-pollinating with limited genetic diversity within each variety but are chosen for their precocity and high rate of bunch emission. Early flowering of the dwarf varieties is a feature of great agronomic relevance that results not only in precocity of nut harvest, but also increased production, while facilitating coconut tree management and harvest operations. While the tall (late) variety takes an average of 8 years to flower, the early flowering dwarfs typically flower on average 2 to 4 years after planting (Perera et al., 2016). The origin of dwarf coconuts has been a matter of debate, although current consensus based on molecular studies points toward a process of domestication from the ancestral tall coconut possibly following the appearance of mutations in flowering time genes resulting in autogamy (Perera et al., 2016), as well gibberellin biosynthesis affecting height growth (Boonkaew et al., 2018; Wang et al., 2021).

Notwithstanding its global socioeconomic importance, advances in coconut tree improvement have been modest, hindered by its perennialism, long generational time, late flowering and large tree size, which demand extensive experimental areas and continuous long-term investments. Genetic improvement programs have adopted two main breeding strategies: (1) intravarietal breeding coupled to mass selection and (2) intervarietal hybridization, the most widely used strategy, which consolidates, into an intermediate F_1_ hybrid, the key features of each variety, such as precocity and short stature from the dwarfs and the rusticity and production of larger fruits from the tall varieties (Batugal et al., 2009; Perera et al., 2016). Rare have been studies in F_2_ populations derived either from selfing or cross pollination among F_1_ Tall x Dwarf hybrids. The wide segregation for production traits reported in F_2_ generations has discouraged such endeavors. Time to flowering, pollination behavior, height and the presence of bole - an expansion at the base of the stem - have been considered to segregate independently. While the former two traits showed the involvement of several genes, height and bole character segregation suggested the presence of at least one major gene each, with incomplete dominance (Fernando and Perera, 1997; Namboothiri et al., 2011; Perera et al., 2016), although no genetic follow-up testing of such hypothesis was ever reported.

Advances in mapping quantitative trait loci (QTLs) of economically important traits have been limited in coconut, restricted to a few studies using Tall x Dwarf F_1_ hybrid populations (Herrán et al., 2000; Ritter et al., 2000; Lebrun et al., 2001; Baudouin et al., 2006; Riedel et al., 2009). In such F_1_ experimental populations, segregating variation can only be captured from the outbred tall coconut, because the dwarf parent, being essentially an inbred line, will not contribute allelic segregation. Flowering time as well as other traits related to the dwarf behavior typically display an intermediate phenotype in the F_1_ progeny with no segregation that would allow genetic mapping of major effects contributed by the dwarf genotype. Studies of F_2_ or backcross populations are needed to advance the molecular understating of key traits related to precocity in coconut. Furthermore, genetic mapping in the cited studies has been carried out either with low portability AFLP markers or a limited set of microsatellites, which do not provide adequate sequence-level information for deeper investigation of putative genes underlying mapped QTLs. Only recently has this scenario begun to change with studies employing single nucleotide polymorphism (SNP) markers for diversity studies (Muñoz-Pérez et al., 2022), linkage mapping to support genome assembly (Yang et al., 2021), and whole genome sequencing for genome-wide studies (GWAS) (Wang et al., 2021). Understanding the genetic basis of morphological traits that determine precocity with a particular interest in early flowering, would represent an important step towards breeding advancement of coconut.

In the present study, we used a unique mapping approach in an outbred F_2_ population generated by open pollination intercrossing F_1_ hybrid trees between the Brazilian Green Dwarf from Jiqui (BGDJ) and the West African Tall (WAT). We exploited the fact that the Dwarf variety is essentially a fixed inbred line with very little if any heterozygosity, while the Tall variety is genetically diverse. Following interbreeding of the F_1_ trees, SNP markers fixed in homozygosity in the Tall variety for the alternative allele to the one fixed in the Dwarf will segregate in a 1:2:1 ratio providing a very large number of loci segregating in a bona fide F_2_ configuration to allow mapping the quantitative variation underlying the genetic divergence between tall and dwarf coconut types. This SNP selection strategy essentially converted the outbred F_2_ population into a standard F_2_, allowing the appropriate application of linkage and QTL mapping. We built a high-density linkage map using SNPs derived from genotyping by sequencing with DArTseq, aligned the map to the current genome assemblies and located QTLs for traits related to precocity, including a major effect locus for flowering time colocalized with the *FLOWERING LOCUS T* (*FT*).

## 2. Materials and methods

### 2.1 Plant material and DNA extraction

The study was carried out with 182 F_2_ plants, generated by open pollination interbreeding among PB-141 F_1_ coconut hybrid plants in a ten-year-old commercial plantation managed by Embrapa Tabuleiros Costeiros, Aracaju – SE (10.9513° South, 37.0528° West). PB-141 is an F_1_ hybrid derived from the cross between the Brazilian Green Dwarf from Jequi (BGDJ) and West African Tall (WAT). Coconuts were collected from F_1_ palms in the central area of the commercial plantation in order to maximize the probability of being effectively derived from intercrossing of F_1_ plants. F_2_ plants were therefore derived either from selfing or cross pollination between different F_1_ plants. Samples of young leaves were collected and used to extract total genomic DNA using an optimized Sorbitol-CTAB-based protocol (Inglis et al., 2018). DNA concentrations and purity were estimated using a Nanodrop 2000 spectrophotometer (Thermo Scientific).

### 2.2 SNP genotyping

Genomic DNA samples were shipped to the Service of Genetic Analysis for Agriculture (SAGA) laboratory in Mexico for high-throughput genotyping using the DArTSeq method developed by Diversity Arrays Technology Pty Ltd. (Sansaloni et al., 2011). DNA complexity reduction was performed using the SbfI/MseI enzyme combination. Samples were processed in digestion/ligation reactions and the scrambled fragments were amplified using two primers with sequence complementary to the ligated adapters and oligonucleotides from the sequencing platform. Subsequently, the clusters were sequenced on an Illumina HiSeq2500 sequencing platform. SNP markers were detected following mapping of sequence reads sampled at adequate depth on the Coconut reference genome (Yang et al., 2021) according to standard parameters defined in DArTsoft14, an automated genomics data analysis program and DArTdb, a laboratory management system developed and patented by DArT for generating SNP marker data. Following the standard scoring system provided by the DArTsoft14 platform, codominant SNP markers were scored as “0” for homozygous reference allele, “1” for homozygous alternative allele, and “2” for presence of both the reference and the alternative allele. These scores were later converted to the corresponding genotypic classes with formats hh, kk and hk respectively, for linkage mapping under an unknown linkage phase configuration (see below).

### 2.3 Morphological and flowering trait evaluations

Due to restrictions in field availability for planting, only 125 of the 182 F_2_ plants were ultimately planted in a field trial in a site known for its soil homogeneity at Embrapa Tabuleiros Costeiros research center in June, 2018. Since no replicate plants were possible for the 125 F_2_ plants (n =125 entries), these were deployed in an augmented randomized complete block design (ARCBD) (Crossa and Federer, 2012) with c = 3 checks (random plants of the parents BGDJ, WAT and the F_1_ hybrid BGDJxWAT), r= 10 blocks extending in rows. Because the number of entries was not a multiple of the number of blocks, nine blocks had 13 entries and the 10^th^ block had 8 entries. Nine traits were ultimately evaluated in 121 live coconut trees: Number of Leaves (NL), Height of Second Leaf Hem (cm) (HSLH), Petiole Length (cm) (PL), Rachis Length (cm) (RL), Petiole Width (cm) (PW), Petiole Thickness (cm) (PT), emergence of the base of the stipe (BOLE), Number of Bunches with fruits (NBF) and Days to Launch of the First Spathe (DLFS). BOLE was recorded in a binary fashion with presence of bole when it extended at least 70 cm from the ground, and absence when at less than 70 cm from the ground. DLFS was used as a direct quantitative measure of time to flowering. These traits, but BOLE, are typically used to differentiate dwarf from tall plants in the early stages of life. Measurements were taken according to the standardized methods described by Santos et al. (1996). Trait evaluations were carried out at one and two years after planting, on June 30, 2019 and July 30, 2020 with evaluation of morphological leaf traits. Flowering initiation was measured on each individual plant according to the emergence of the first spathe starting on October 11, 2020 and then by a monthly monitoring to precisely catch the launch of the first spathe. The number of days for the emergence and opening of the subsequent spathes until the tenth spathe was recorded until the least evaluation on August 11, 2022. The final evaluations of the morphological traits were carried out on May 10, 2022, with the exception of NBF, which was evaluated on August 11, 2022. Pearson correlations were estimated to assess the correlation between traits using the Corrplot package available in R (Wei et al., 2021). Phenotypic data for all traits but BOLE was analyzed with the following mixed model: *Y_ij_ = μ + B_i_ + E_j_ + C (_i_)_j_ + e_ij_* where *Y* is the observed phenotypic value of the genotype entry (*j)* in block (*i*), *μ* is the overall mean, *B* is the fixed block effect, *E* is the random genotype entry effect, *C* is the random effect of the common checks in blocks and *e* is the random residual effect. Best linear unbiased predictors (BLUPs) were estimated using the software SELEGEN (Resende, 2016) under model 76 (augmented blocks, genotypes, one plant per plot, single location, fixed check).

### 2.4 Linkage map construction

The raw SNP calls were filtered for quality for the linkage mapping analyses. The first filtering step was based on a call rate >0.95 followed by selection of SNP markers that fit the expected segregation proportions in an F_2_, by a Chi-square test under the null hypothesis of a 1:2:1 segregation ratio with a slightly more relaxed threshold (*α*>0.001) to allow including SNPs even with some level of segregation distortion given the particularity of the mapping population. Linkage analysis and construction of the genetic map were performed according to the standard F_2_ model using JoinMap v3.0 (Van Ooijen and Voorrips, 2001). The genotypes of the F_2_ population were coded, according to the format <hkxhk> for an unknown linkage phase configuration and three possible genotypic classes. Markers were assigned to linkage groups (LG) via the clustering module with LOD (logarithm of the odds) scores greater than 15, maximum recombination fraction of 0.4, under a Kosambi mapping function. Linkage maps were drawn and aligned to the assembled genomes using MAPCHART (Voorrips, 2002). For the alignments, recombination distances were expressed in centiMorgans and physical distances in Mbp.

### 2.5 Marey maps and recombination rates in the genome of *Cocos nucifera*

To estimate local recombination rates across the 16 coconut chromosomes, the 69 base pair DArTseq sequences corresponding to the SNP markers were aligned to the two existing chromosome-level coconut genome assemblies of Hainan Tall coconut: (1) a draft assembly built using exclusively short Illumina paired-end sequencing (Yang et al., 2021) hereafter called the YANG genome and (2) a reference-grade assembled genome using long Nanopore single-molecule sequencing and Hi-C technology (Wang et al., 2021) hereafter called the WANG genome. The FASTA sequences of the chromosome assemblies were downloaded from their respective repositories. The YANG genome from GenBank BIOPROJECT PRJNA374600, and the WANG genome from the Genome Warehouse in the National Genomics Data Center (Beijing Institute of Genomics) accession number GWHBEBT00000000, BIOPROJECT PRJCA005463. The DArTseq sequences were mapped to the sequences of the 16 coconut pseudo-chromosomes using BLASTn for highly similar sequences (megablast) with a cutoff E-value <1E-05. DArTSeq sequences that mapped to a unique location in both genome versions were used in further analyses. The relationships between the genetic and physical positions of the mapped SNPs were represented by the local slope of the curve adjusted to the datapoints in MAREY maps (Chakravarti, 1991). Curves were fitted to the data using locally weighted scatterplot smoothing (LOESS) polynomial regression with a smoothing parameter of 0.3 using MareyMap Online (Siberchicot et al., 2017). Sporadic abnormal points from the map that did not fit the expected monotonously increasing relationship between physical and genetic positions were removed using the available functionality in the software. The physically mapped markers were used to obtain an estimate of the effective physical coverage achieved by the genetic map and the recombination rates in cM/Mb along the chromosomes. Pearson’s correlations along windows of 1 Mb were calculated between the average recombination rate in the 1 Mb interval and the number of genes as annotated in the coconut chromosomes in the YANG genome (Yang et al., 2021).

### 2.6 QTL mapping

QTLs detection was performed using the BLUPs for eight traits and the binary 0/1 value for BOLE with R/qtl version 3.5.3, using three different methods: interval mapping (IM), Haley Knott regression (HK) and non-parametric interval mapping (Non-parametric IM) for those traits that violated the assumption of normality (Broman et al., 2003). The probability of genotype error (error.prob = 0.01) was calculated and the threshold LOD score for QTL detection was determined by genome-wide LOD significance thresholds, setting LOD thresholds at different alpha levels (α = 0.01; 0.05 and 0.1) using 1900 permutations to allow for the declaration of QTLs with variable stringency levels in order to control also for Type II errors.

### 2.7 Co-localization of the FLOWERING LOCUS T (FT) gene with the flowering time QTL

A set of 198 expressed gene sequences in coconut were recently identified as homologues of flowering-time genes in *Arabidopsis*. Five of them, annotated as FT homologs to the *Arabidopsis thaliana* FT gene (AT1G65480) were found to be differentially expressed between seedling and reproductive leaf samples in both tall and dwarf coconut. Alternative splicing analysis showed that the FT gene produces different transcripts in tall compared to dwarf coconut (Xia et al., 2020). We took all the 87 *Arabidopsis thaliana* genes present in the annotation of the 198 coconut expressed genes and aligned them to the QTL region on the corresponding chromosomes assembled in the two genome assemblies using BLASTn specifically in the genome stretch from 65 Mbp to 75 Mbp where the major effect QTL was mapped (see below).

## 3. Results

### 3.1 SNP markers and linkage mapping

Genotyping by sequencing using DArTseq generated 86,719 raw SNP calls based on the standard parameters used in DArTsoft. After applying a call rate threshold of 95% and a test for adherence to a null hypothesis of 1:2:1 segregation, 3,714 SNPs (4.28% of the raw SNP calls) were retained for further linkage analyses (File S1). Dominant presence/absence in silico DArTseq variants were not used due to their low information content and considering that an adequate number of co-dominant informative SNPs was already available for linkage mapping purposes. A linkage map was ultimately built with 182 F_2_ individuals that correspond to 364 meiotic events sampled in the gametes from the recombined F_1_ parents. All 3,714 SNPs were mapped with LOD ≥15 and maximum recombination fraction of 0.4, in 16 linkage groups matching the expected 16 coconut chromosomes. However, for the subsequent assessment of the recombination rates and QTL mapping, only the SNPs that were physically positioned on the genome assemblies (see below) were ultimately considered on the linkage map (File S2). Linkage group numbering used for our map was based on the physical positioning of SNP markers on the YANG genome chromosomes which was published first in the literature. The chromosome numbering used in the WANG genome was different but the correspondence was established and reported throughout (Table 1). An average of 184.5 SNPs were mapped to each chromosome, varying from a maximum of 329 SNPs on chromosome 1 to a minimum of 101 SNPs on chromosome 15. The genetic map spanned a total of 2,124.3 cM in recombination size, with an average linkage map length of 132.8 cM, and an average intermarker recombination distance between adjacent marker pairs of 0.742 cM. Chromosome 15 displayed a large recombination gap of 31.2 cM between two denser linkage clusters although still linked by the minimum LOD 15 score threshold adopted (Figure 1). Except for this large gap observed on chromosome 15, maximum intermarker distance was 6.7 cM on chromosome 7.

**Figure 1.**
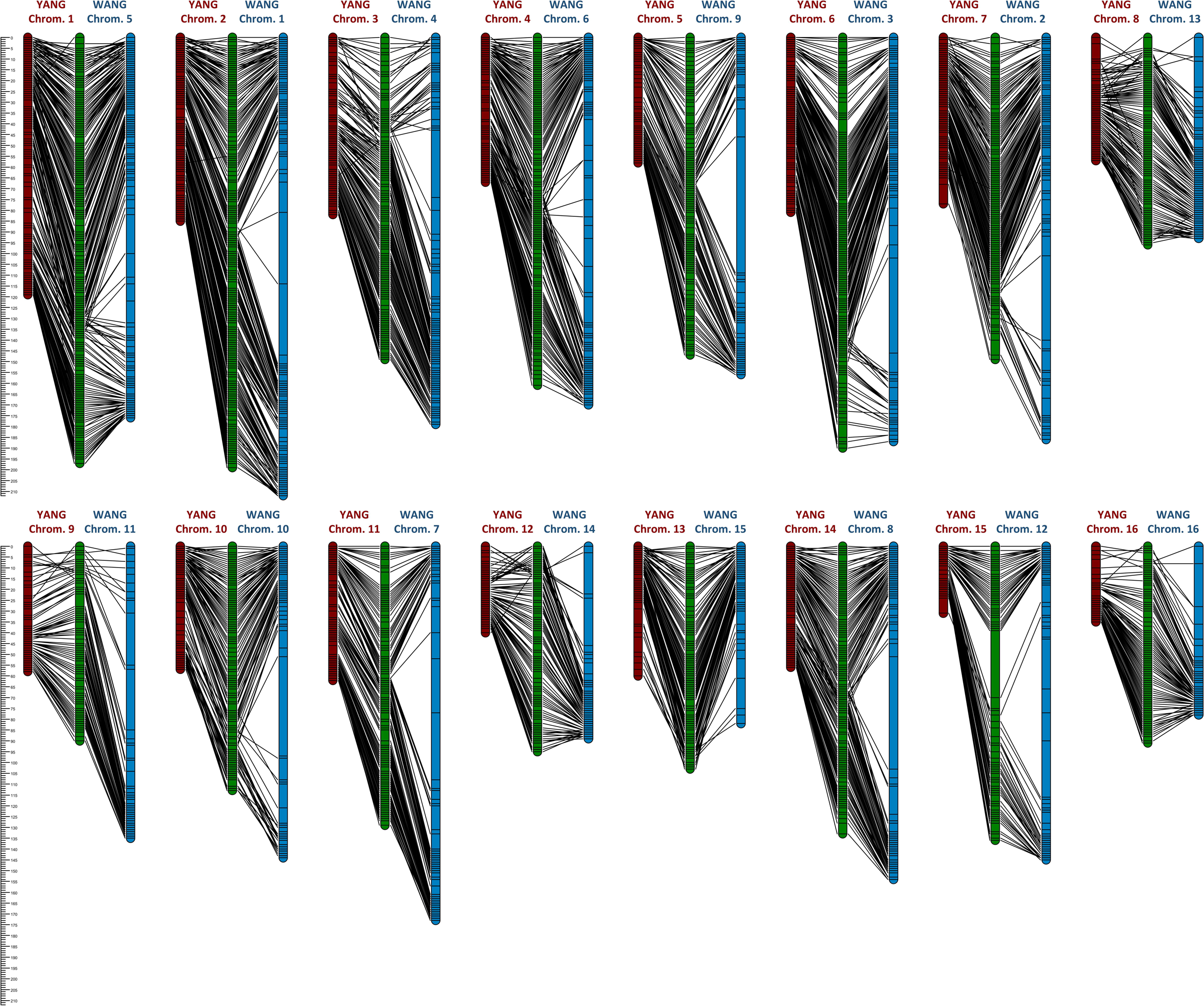
Alignment of the linkage map (green) to the two chromosome-level genome assemblies, the YANG genome assembly (red) (Yang et al., 2021) and the WANG genome assembly (blue) (Wang et al., 2021).

**Table 1.**
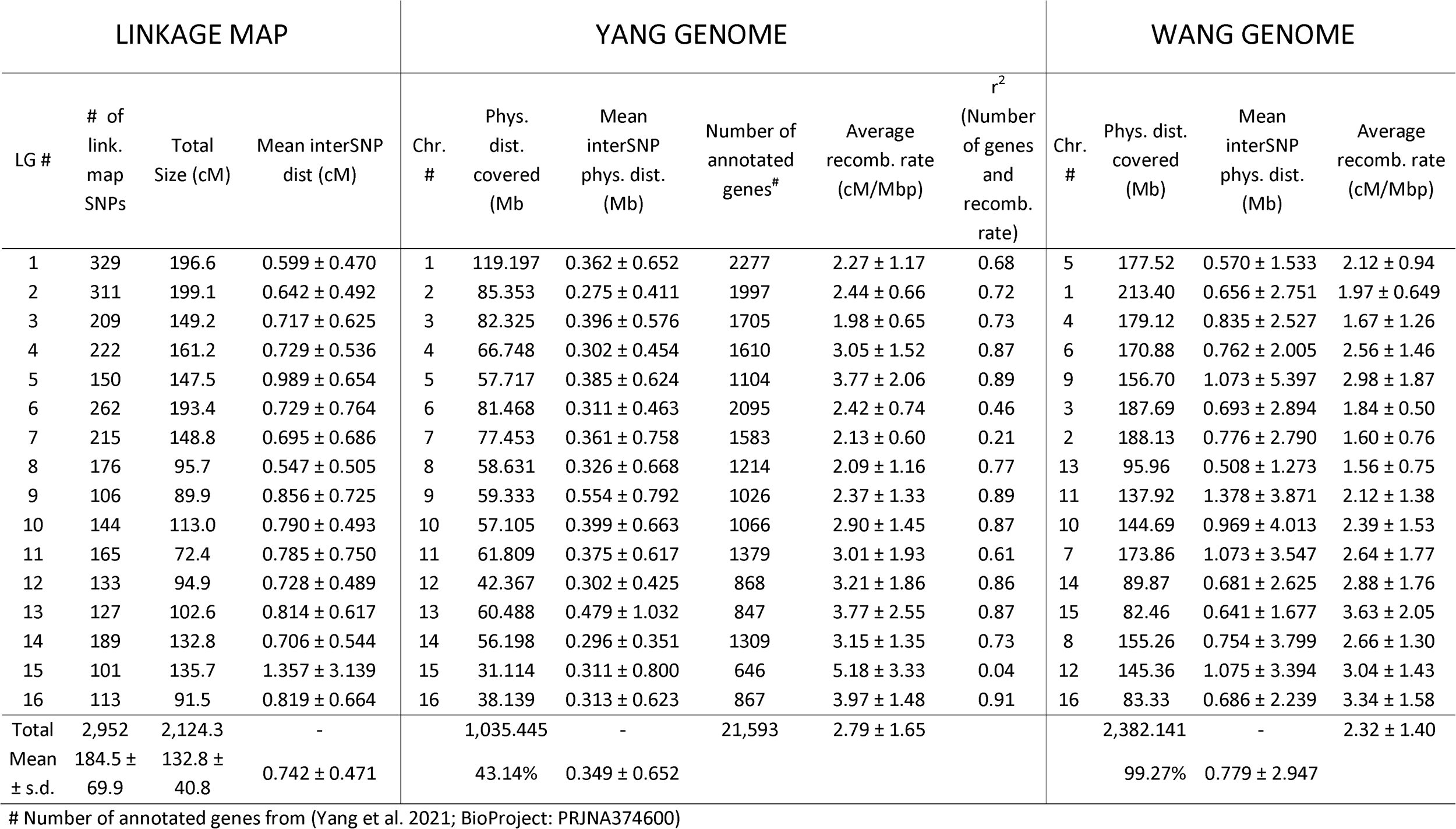
Statistics of the linkage map and its alignment to the two available chromosome-level genome assemblies of coconut (Cocos nucifera L.).

| LINKAGE MAP |  |  |  | YANG GENOME |  |  |  |  |  | WANG GENOME |  |  |  |
| --- | --- | --- | --- | --- | --- | --- | --- | --- | --- | --- | --- | --- | --- |
| LG # | # of link. map SNPs | Total Size (cM) | Mean interSNP dist (cM) | Chr. # | Phys. dist. covered (Mb) | Mean interSNP phys. dist. (Mb) | Number of annotated genes<sup>#</sup> | Average recomb. rate (cM/Mbp) | r<sup>2</sup> (Number of genes and recomb. rate) | Chr. # | Phys. dist. covered (Mb) | Mean interSNP phys. dist. (Mb) | Average recomb. rate (cM/Mbp) |
| 1 | 329 | 196.6 | 0.599 ± 0.470 | 1 | 119.197 | 0.362 ± 0.652 | 2277 | 2.27 ± 1.17 | 0.68 | 5 | 177.52 | 0.570 ± 1.533 | 2.12 ± 0.94 |
| 2 | 311 | 199.1 | 0.642 ± 0.492 | 2 | 85.353 | 0.275 ± 0.411 | 1997 | 2.44 ± 0.66 | 0.72 | 1 | 213.40 | 0.656 ± 2.751 | 1.97 ± 0.649 |
| 3 | 209 | 149.2 | 0.717 ± 0.625 | 3 | 82.325 | 0.396 ± 0.576 | 1705 | 1.98 ± 0.65 | 0.73 | 4 | 179.12 | 0.835 ± 2.527 | 1.67 ± 1.26 |
| 4 | 222 | 161.2 | 0.729 ± 0.536 | 4 | 66.748 | 0.302 ± 0.454 | 1610 | 3.05 ± 1.52 | 0.87 | 6 | 170.88 | 0.762 ± 2.005 | 2.56 ± 1.46 |
| 5 | 150 | 147.5 | 0.989 ± 0.654 | 5 | 57.717 | 0.385 ± 0.624 | 1104 | 3.77 ± 2.06 | 0.89 | 9 | 156.70 | 1.073 ± 5.397 | 2.98 ± 1.87 |
| 6 | 262 | 193.4 | 0.729 ± 0.764 | 6 | 81.468 | 0.311 ± 0.463 | 2095 | 2.42 ± 0.74 | 0.46 | 3 | 187.69 | 0.693 ± 2.894 | 1.84 ± 0.50 |
| 7 | 215 | 148.8 | 0.695 ± 0.686 | 7 | 77.453 | 0.361 ± 0.758 | 1583 | 2.13 ± 0.60 | 0.21 | 2 | 188.13 | 0.776 ± 2.790 | 1.60 ± 0.76 |
| 8 | 176 | 95.7 | 0.547 ± 0.505 | 8 | 58.631 | 0.326 ± 0.668 | 1214 | 2.09 ± 1.16 | 0.77 | 13 | 95.96 | 0.508 ± 1.273 | 1.56 ± 0.75 |
| 9 | 106 | 89.9 | 0.856 ± 0.725 | 9 | 59.333 | 0.554 ± 0.792 | 1026 | 2.37 ± 1.33 | 0.89 | 11 | 137.92 | 1.378 ± 3.871 | 2.12 ± 1.38 |
| 10 | 144 | 113.0 | 0.790 ± 0.493 | 10 | 57.105 | 0.399 ± 0.663 | 1066 | 2.90 ± 1.45 | 0.87 | 10 | 144.69 | 0.969 ± 4.013 | 2.39 ± 1.53 |
| 11 | 165 | 72.4 | 0.785 ± 0.750 | 11 | 61.809 | 0.375 ± 0.617 | 1379 | 3.01 ± 1.93 | 0.61 | 7 | 173.86 | 1.073 ± 3.547 | 2.64 ± 1.77 |
| 12 | 133 | 94.9 | 0.728 ± 0.489 | 12 | 42.367 | 0.302 ± 0.425 | 868 | 3.21 ± 1.86 | 0.86 | 14 | 89.87 | 0.681 ± 2.625 | 2.88 ± 1.76 |
| 13 | 127 | 102.6 | 0.814 ± 0.617 | 13 | 60.488 | 0.479 ± 1.032 | 847 | 3.77 ± 2.55 | 0.87 | 15 | 82.46 | 0.641 ± 1.677 | 3.63 ± 2.05 |
| 14 | 189 | 132.8 | 0.706 ± 0.544 | 14 | 56.198 | 0.296 ± 0.351 | 1309 | 3.15 ± 1.35 | 0.73 | 8 | 155.26 | 0.754 ± 3.799 | 2.66 ± 1.30 |
| 15 | 101 | 135.7 | 1.357 ± 3.139 | 15 | 31.114 | 0.311 ± 0.800 | 646 | 5.18 ± 3.33 | 0.04 | 12 | 145.36 | 1.075 ± 3.394 | 3.04 ± 1.43 |
| 16 | 113 | 91.5 | 0.819 ± 0.664 | 16 | 38.139 | 0.313 ± 0.623 | 867 | 3.97 ± 1.48 | 0.91 | 16 | 83.33 | 0.686 ± 2.239 | 3.34 ± 1.58 |
| Total | 2,952 | 2,124.3 | - |  | 1,035.445 | - | 21,593 | 2.79 ± 1.65 |  |  | 2,382.141 | - | 2.32 ± 1.40 |
| Mean ± s.d. | 184.5 ± 69.9 | 132.8 ± 40.8 | 0.742 ± 0.471 |  | 43.14% | 0.349 ± 0.652 |  |  |  |  | 99.27% | 0.779 ± 2.947 |  |

### 3.2 Alignment of the linkage map to the genome assembly

The 3,714 DArTseq sequences containing the linkage mapped SNPs were submitted to BLASTn against the chromosome assemblies. On the YANG genome out of the 3,714 linkage mapped DArTSeq SNPs, 2,952 DArTSeq sequences were mapped to unique locations with E-values <7.3e-05. The sequences for the remaining SNPs either aligned to multiple locations or did not align to the assembled coconut chromosomes and were removed from further analyses. On the WANG genome 3,027 DArTSeq sequences were mapped to unique locations with E-values <3.6e-11 (File S3). On the YANG genome, the linkage map covered 1,035 Mb corresponding to a coverage of approximately 43.14% of the 2,400 Mb *Cocos nucifera* Tall cultivar genome (Wang et al., 2021). On the WANG genome, the linkage mapped SNPs covered a total of 2,382 Mb, a 99.27% coverage, more than twice the coverage obtained on the YANG genome with a corresponding lower density of mapped SNPs and long stretches with no segregating SNPs mapped to them. The average estimated physical inter SNP distance were 349kb and 779kb, SNP density of 2.85 and 1.27 SNPs/Mb and maximum physical distances of 9.7 and 63.4 Mb respectively on the YANG and WANG genomes (Table 1).

### 3.3 Marey maps and recombination rates

The linkage maps alignment to the 16 pseudochromosomes was largely colinear with the physical assembly, with only sporadic inconsistencies of the map versus genome positions, 2 in the YANG genome and 20 in the WANG genome (Figure 1; File S4). As expected, on all linkage groups the alignment to the physically larger WANG genome showed long sections of genome sequence with few or no SNP markers mapped. The Marey maps showed plateaus of restricted recombination, where the recombination distance does not follow the increase in physical distance along the chromosome (Figure 2). These plateaus spanned considerably longer stretches for the Marey maps on the WANG genome. From the Marey maps data, estimates of local recombination rate were obtained for both genomes using LOESS (Locally Weighted Scatterplot Smoothing) (File S4) and plotted for the YANG genome (Figure 2). Following LOESS treatment of the data, the average local recombination rates were similar: 2.78 cM/Mb for the YANG genome and 2.32 cM/Mb for the WANG genome. The smoother recombination rate curves closely followed expectations. Chromosomal segments of restricted recombination in the Marey maps coincided with local valleys of low recombination rate notably spotted on YANG chromosomes 1, 4, 5, 11 and 14, for example and more accentuated on the corresponding WANG chromosomes.

**Figure 2.**
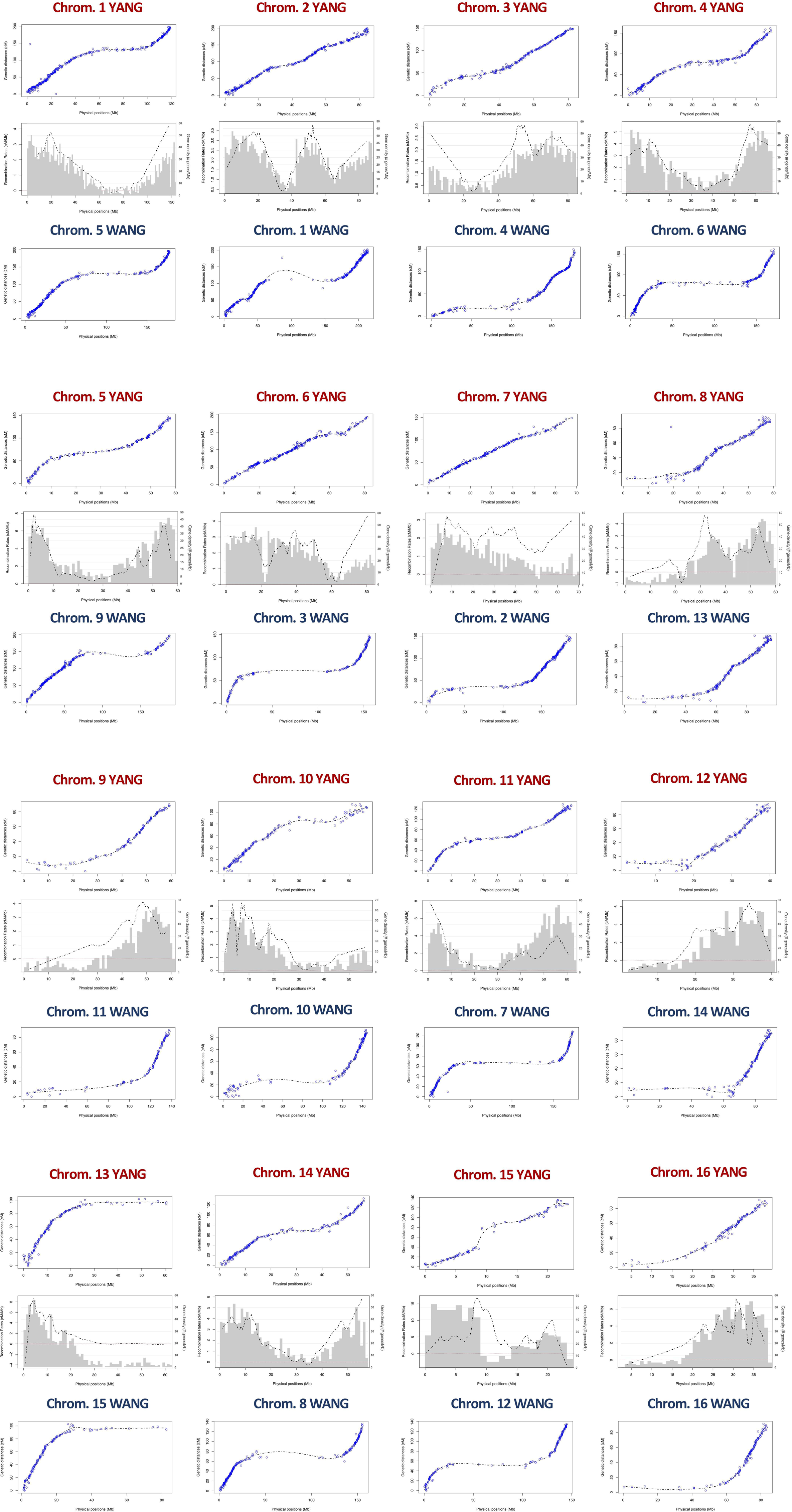
Marey maps showing the correspondence between the physical position (x-axis, Mb) and the recombination-based position (y-axis, cM) on the 16 Coconut chromosomes of the YANG genome assembly (Yang et al., 2021) and the WANG genome assembly (Wang et al., 2021). Also shown in the intermediate panels the combined pattern of gene density distribution and the LOESS curve of recombination rate for the YANG genome.

Average recombination rates per chromosomes varied across chromosomes up to more than twofold (e.g., 1.98 cM/Mb on chromosome 3 and 5.18 cm/Mb on chromosome 15 on the YANG genome). However, the correlation between the rates estimated on the two genome versions was high and significant (r = 0.85; p-value = 3E-5). The highest rate on chromosome 15 is likely the result of the large linkage gap observed on this chromosome. The average recombination rate across the entire genome were similar, at 2.79 and 2.32 cM/Mb for the two genomes (Table 1). Furthermore, taking advantage of the gene annotation provided for the YANG genome sequence, we underlaid the frequency distribution of the number of genes in 1 Mb intervals, equivalent to gene density, to the smoothed recombination rate curve in the same 1 Mb segment (Figure 2). Visually an almost perfect direct relationship was seen between gene density and recombination rate. This visual relationship was further validated by observing high and significant Pearson correlation between average recombination rate and number of genes in the 1 Mb intervals estimated for the YANG genome. For all chromosomes but chromosomes 7 and 15 these correlations were high and significant (p-value <0.001) (Table 1) suggesting, once again, some different behavior as far as recombination for these two chromosomes.

### 3.4 Trait variation and correlations

The F_2_ individuals displayed extensive segregation for all traits measured, as expected in a recombinant F_2_ population of divergent cultivars. Raw phenotypic values and BLUPs are provided in File S5. The continuous traits showed an approximately normal distribution, although with some skewness, most notably for NBF and DLFS (Figure 3). BOLE expression fit a 3:1 segregation ratio with 98 plants showing no bole and 23 with bole (Chi-square 2.31; p-value = 0.128). Days to Launch of the First Spathe (DLFS) varied from a minimum of 812 days for the precocious flowering plants to a maximum of 1608 days for the late blooming ones with an average of 1287 ± 214.5 days. One plant therefore displayed 2.2 standard deviations below the mean and 13 flowered with less than 1000 days, corresponding to more than one standard deviation below the mean. Eight of these very early flowering plants displayed the highest numbers (≥13) fruit per bunch (NBF). Significant negative correlations were seen between DLFS and NL, HSLH, BOLE and NBF, and a significant positive correlation with PL (Figure 4). Positive significant correlations were seen between five of the six leaf morphology traits (NL, HSHL, RL, PW and PT), except PL that showed no correlation with all other five traits. Positive and significant correlations were also observed between NBF and NL, HSLH, RL and BOLE. Coconut trees with a recorded bole flowered on average after 1098 days and produced an average of 9.5 fruit per bunch while plants without bole flowered on average after 1347 days, and produced 5.96 fruit per bunch.

**Figure 3.**
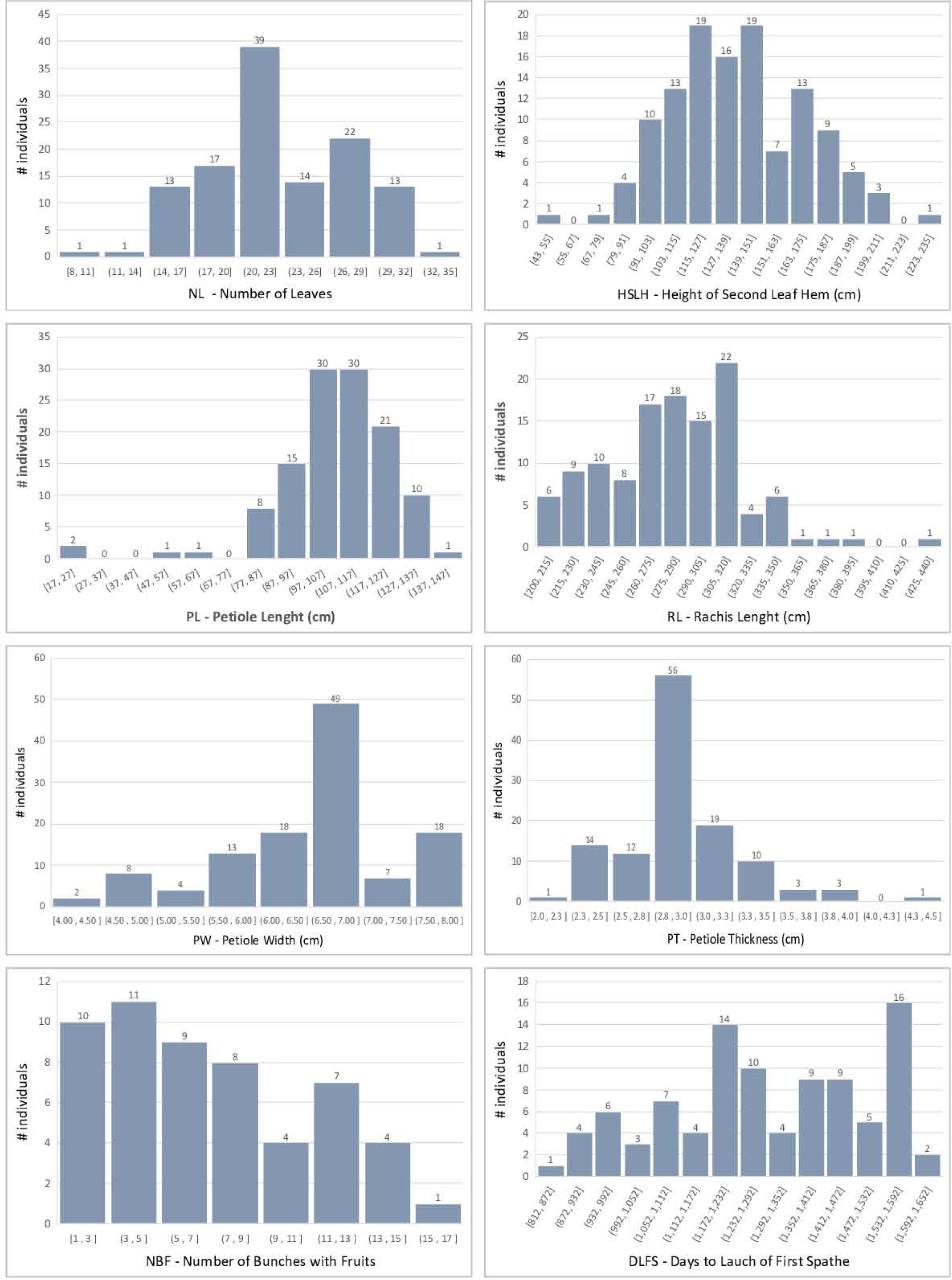
Histogram of the coconut phenotypic trait distributions in the F_2_ mapping population.

**Figure 4.**
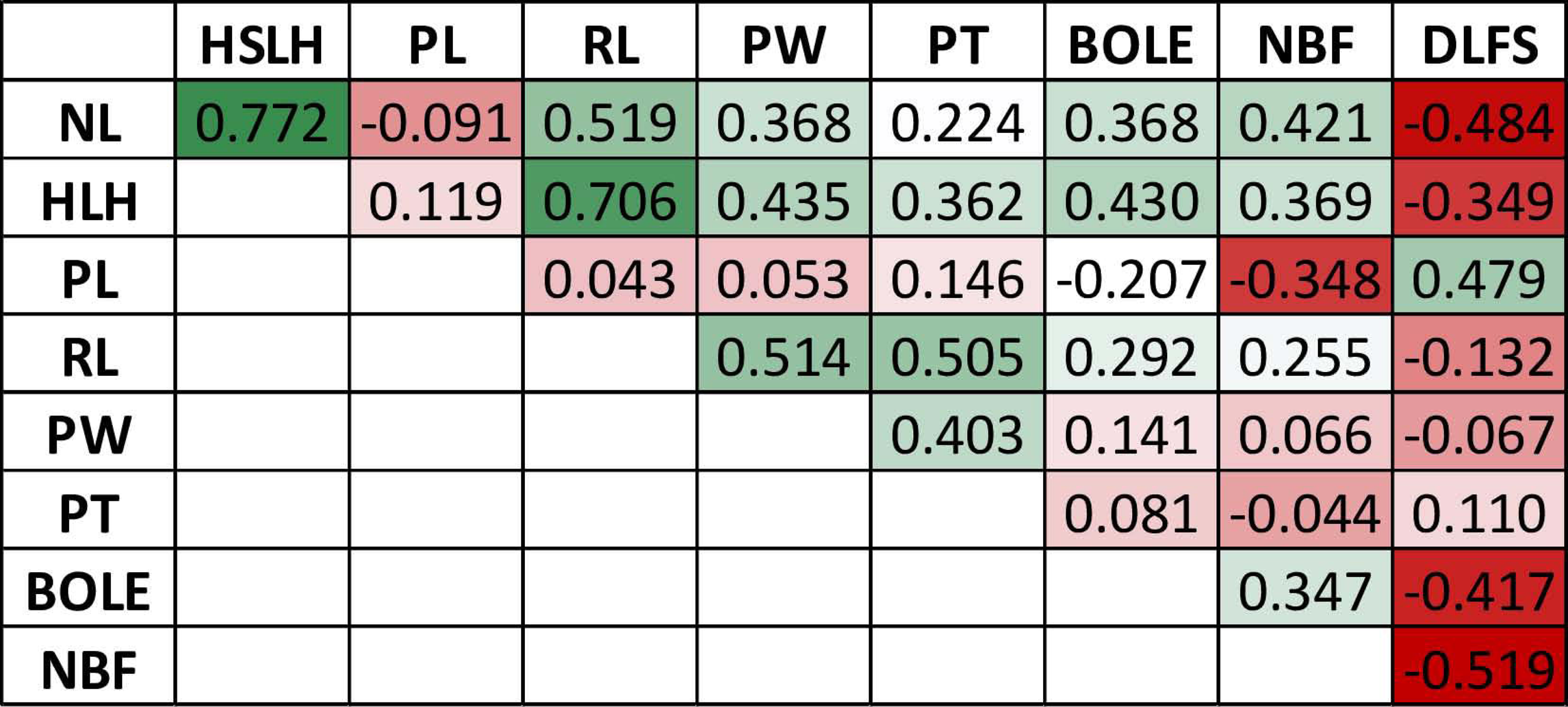
Matrix of genotypic correlations among the measured traits in the F_2_ mapping population

### 3.5 Mapping and haplotype analysis of a major flowering time QTL

QTL detection was carried out with three methods, IM, CIM and HK to assess consistency of results and minimize type I errors. Significant QTL peaks were declared based on the LOD limit analysis for the different alpha levels to allow the detection of suggestive QTLs in order to minimize Type II errors. The significance thresholds indicated LOD limits of 4.15, 4.49 and 5.22 respectively for the different alpha levels (α = 0.01; 0.05 and 0.1) (Table 2). Results with CIM showed a considerably larger number of 16 putatively significant QTLs in 11 chromosomes when compared to the results with IM and HK. These CIM results are not discussed further but are provided (File S6). Genome-wide QTL profile results across the 16 chromosomes obtained with the three QTL detection methods, for the four traits for which significant QTLs were detected are provided for a full visualization of results. The LOD limit thresholds are specified by the differentially dotted lines (Figure 5).

**Figure 5.**
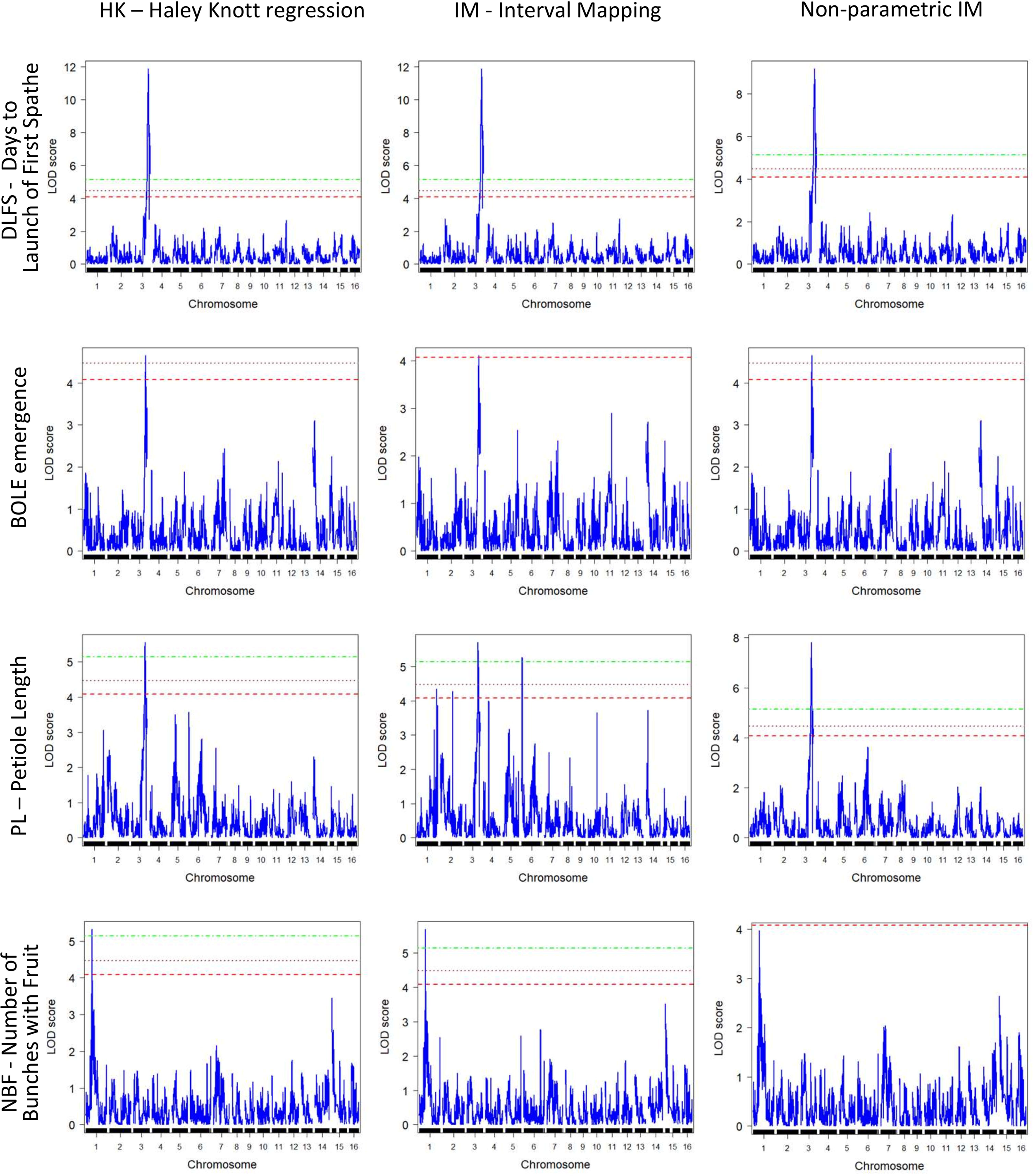
Genome-wide QTL mapping profiles for the mapped traits (Y axis) obtained with the three different QTL mapping methods implemented (X axis)

**Table 2.** Summary of QTL mapping results in the coconut F_2_ population.

|  | **Days to Launch of**<br>**First Spathe (DLFS)** | **Bole emergence**<br>**(BOLE)** | **Number of bunches**<br>**with fruit (NBF)** | **Petiole Length (PL)** |
| --- | --- | --- | --- | --- |
| **Chromosome** | 3 | 3 | 1 | 3 |
| **LOD score threshold**<br>**(percentile)** | 5.14 (99%) | 4.47 (95%) | 5.14 (99%) | 5.14 (99%) |
| **# SNPs in QTL region** | 52 | 1 | 1 | 22 |
| **QTL region position (cM)** | 118.928 - 149.166 | 132.62 | 57.16 | 123.503 – 136.125 |
| **Right SNP** | 100287206_T>A-55 | - | - | 100287206_T>A-55 |
| **Left SNP** | 100273632_T>G-15 | - | - | 100273632_T>G-15 |
| **Peak LOD** | 11.86 | 4.64 | 5.68 | 7.79 |
| **Peak LOD SNP** | 100226499_C>T-34 | 100228320_G>A-40 | 100237259_A>G-48 | 100226499_C>T-34 |
| **Target SNP allele** | T | A | A | T |
| **(Origin; effect)** | (Dwarf; reduction) | (Dwarf; increase) | (Dwarf; increase) | (Dwarf; reduction) |
| **Genome position (bp)** | 73,147,443 | 75,613,473 | 19,206,916 | 73,147,443 |
| **Variance explained (%)** | 64.0 | 3.12 | 0.43 | 22.3 |

Overall, consistent QTLs across the methods were detected on chromosome 3 for DFLS, BOLE and PL and on chromosome 1 for NBF. A genomic segment at the tip of chromosome 3 spanning 52 significantly linked SNPs to DFLS stands out as a major effect QTL for flowering time, explaining an estimated 64% of the variation in DLFS. The alignment of the linkage map to the chromosome 3 of the YANG genome and chromosome 4 of the WANG genome allowed a detailed haplotype analysis of this segment. In the YANG genome starts at SNP 100285601_C>G-14 with LOD 4.99 located at 117.99 cM corresponding to 68.84 Mbp, and ends at SNP 100270789_A>G-59 with LOD 6.53 located at 148.17 cM corresponding to position 82.32 Mbp (File S3). In the WANG genome it spans 55 SNPs including the 52 mapped to the YANG genome form position 162.7 Mb to the end of the chromosome at 178.5 Mb. Within this segment, the highest QTL peak (LOD 11.86) is located at SNP 100226499_C>T-34, positioned at 129.11 cM and 73.15 Mbp, flanked by SNPs 100287206_T>A-55 and 100273632_T>G-15. At SNP 100226499_C>T the T allele inherited from the Dwarf parent AVeBrJ causes a major reduction in the time to flower, suggesting incomplete dominance of the locus. Heterozygous genotype G/T confers a reduction from 1460 to 1390 days to the launch of the first spathe, while the homozygous T/T genotype confers a further reduction of almost 300 days, as the coconut trees bearing this genotype flowered on average after 1110 days (Figure 6). In addition to acting on early flowering, the same genomic segment is involved in the control of BOLE and PL. The LOD peak for both QTLs for BOLE and PL was located at the same SNP 100228320_G>A-40, at 132.62 cM and 75.61 Mbp. The co-localization of QTLs for three traits suggests a pleiotropic action of this major effect QTL segment on chromosome 3. Some additional suggestive QTLs were detected for PL on chromosomes 1, 2, 4, 6 that were significant at α = 0.1 based on the IM analysis but that did not reach significance with the two other QTL detection methods. Finally, a single QTL was also found for NBF on chromosome 1 whose Dwarf allele confers an increase in the number of fruits per bunch.

**Figure 6.**
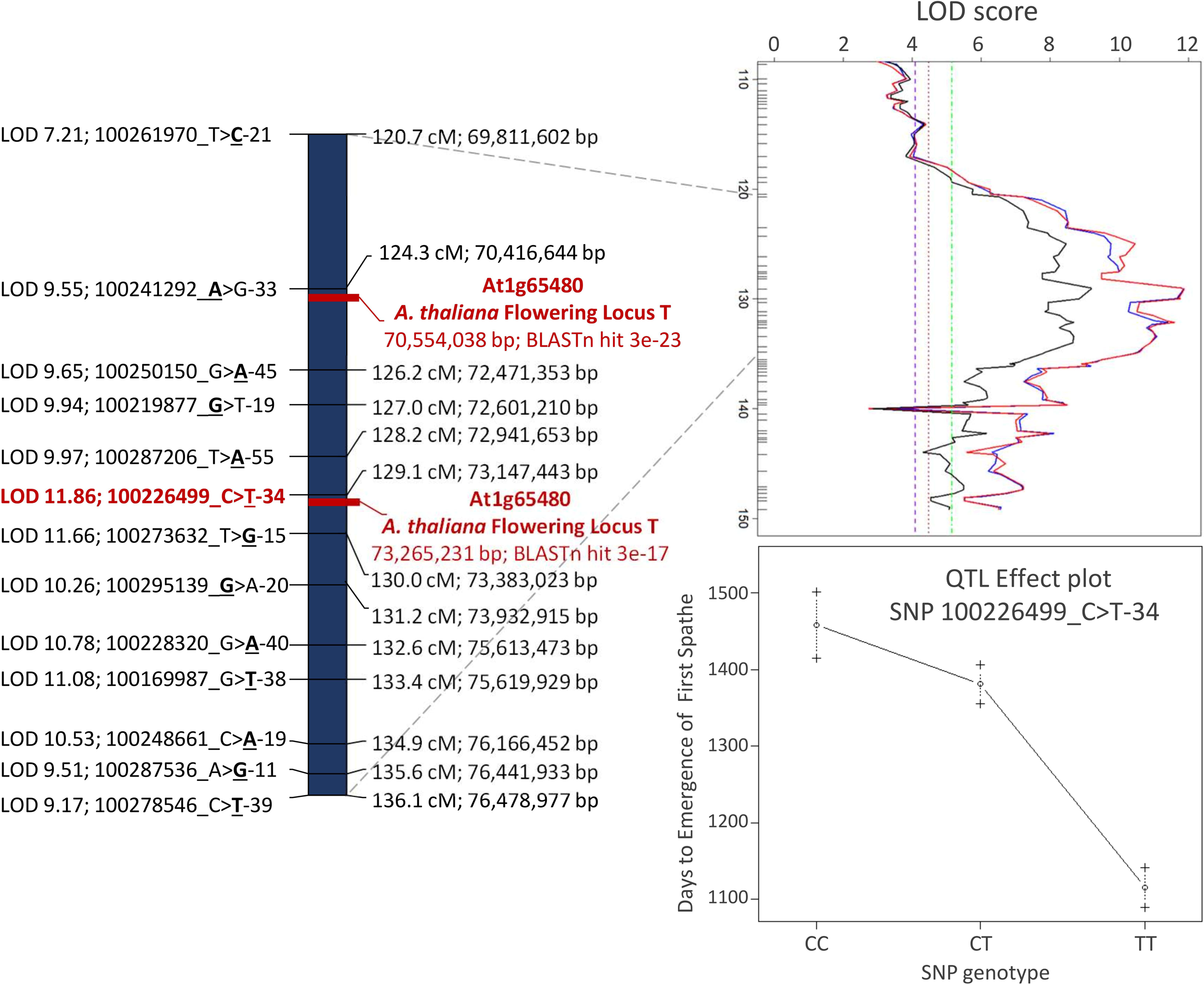
Detailed view of the major effect QTL region for flowering time, petiole length and bole presence in the stem on chromosome 3 of the YANG genome. The *A. thaliana* FLOWERING LOCUS T was colocalized with the leading SNPs for the QTL.

### 3.6 Co-localization of the flowering time QTL with the FLOWERING LOCUS T (FT)

Out of the 87 *Arabidopsis thaliana* genes involved in flowering-time pathways that were found to be homologous to 198 coconut genes (Xia et al., 2020), only the complete coding sequence of the *Arabidopsis thaliana* putative flowering signals mediating protein FT (At1g65480) produced strongly significant alignment to the flowering time QTL mapped on linkage group 3. The FT gene (At1g65480) complete cds mRNA (accession AY065378.1) with 840 bp produced two hits in the YANG genome: (i) at 70,554,038 to 70,554,249 bp with bit score 108, 71% identity, e-value = 3e-23 and 0/212 gaps (0%) and (ii) at 73,265,231 to 73,265,450 bp with bit score 88.7, 70% identity, e-value = 3e-17 and 3/222 (1%) gaps (Figure 6; File S7). These hits are located respectively at 137 kb from SNP 100241292_A>G-33 LOD 9.55 and 117 kb from SNP 100226499_C>T-34 this last one had the highest LOD score (LOD 11.86) in the QTL region. When the FT gene (At1g65480) was blasted against the corresponding chromosome 4 in the WANG genome it also produced two significant hits at 164.688 Mb (e-value = 7E-21) and 168.065 Mb (e-value = 7E-15) that correspond to the genomic positions at the tip of the chromosome (Figure 1; File S8), further corroborating the finding. This co-localization result between the flowering time QTL and the FLOWERING LOCUS T (FT) gene when taken together with the splice variant analysis reported earlier (Xia et al., 2020), strongly suggests that the underlying candidate gene to the flowering time QTL mapped in this study could be an FT homolog in *Cocos nucifera*.

## 4. Discussion

We have carried out QTL mapping in *Cocos nucifera* based on a novel mapping approach for the crop, using an outbred F_2_ population derived from open pollination intercrossing of F_1_ hybrids. With this approach we were able to segregate and track the variation contributed by the inbred Dwarf in the F_2_ by selecting SNP markers fixed for alternative alleles between the Brazilian Green Dwarf from Jiqui (BGDJ) and the West African Tall (WAT) cultivar segregating in a bona fide F_2_ configuration despite the outbred nature of the population. Almost 3,000 SNPs obtained by genotyping by sequencing were positioned on the two existing chromosome-level coconut genome assemblies. By aligning the linkage map to the genome sequences, we provided an independent assessment of the contiguity of the current coconut reference genomes, and a detailed analysis of the relationship between recombination rate and physical distance for the corresponding coconut chromosomes. Furthermore, based on the segregation observed for traits related to precocious behavior inherited from the Dwarf cultivar, we mapped a major effect QTL for flowering time on chromosome 3. Flowering Locus T (FT) gene homologs of coconut, previously described as involved in flowering time by alternative splice variant analysis, were co-localized with the molecular phenotype mapped QTL, providing evidence for its role as the putative underlying gene.

### 4.1 Exploiting the unique genetics of the Tall x Dwarf coconut hybrid for linkage mapping in an outbred F_2_

The linkage map reported in our study is the first one in coconut built from an F_2_ segregating population from a Tall x Dwarf hybrid. Previous maps either used hybrid F_1_ or backcross (BC1) populations (Herrán et al., 2000; Ritter et al., 2000; Lebrun et al., 2001; Yang et al., 2021) or a cross between Tall cultivars (Riedel et al., 2009). Differently from a conventional F_2_ populations generated by selfing an individual F_1_ plant, the particular genetics of coconut allowed using an open pollinated progeny of an F_1_ hybrid field. The parental F_1_ plants composed a commercial production plantation with seeds resulting from controlled pollination between a pollen mix of several Tall coconut parents on female flowers of several Dwarf plants. In the particular case of coconut, we took advantage of the fact that the Dwarf variety is essentially a fixed inbred line with very little if any residual heterozygosity, while the Tall parents are allogamous, heterozygous and genetically diverse (Santos et al., 2020; Wang et al., 2021; Muñoz-Pérez et al., 2022). The F_1_ plants SNP genotypes would potentially differ only in what was inherited from the Tall parents, but SNP markers fixed in the Tall variety and polymorphic in relation to the Dwarf would segregate in a 1:2:1 ratio in the F_2_, irrespective of what Tall parent generated each different F_1_ plant which in turn were intercrossed to produce the F_2_ population. This premise proved valid, and 3,714 SNPs called with a call rate >95%, fit the expected segregation and were linkage mapped. This SNP selection strategy essentially converted the outbred F_2_ population into a standard F_2_, allowing the appropriate application of linkage and QTL mapping. Approximately 20% of the sequences of linkage mapped DArTSeq mapped to multiple locations in the two genomes and were left out of subsequent analysis, as no objective criteria allowed choosing one location over another. Therefore, for QTL mapping analyses we used only around 3,000 linkage mapped SNPs that could be anchored to unique positions on the 16 assembled chromosomes of the two genome versions.

The linkage map assembled in the 16 expected linkage groups spanned a total of 2,124.3 cM, slightly smaller than the linkage map reported earlier by Yang et al. (Yang et al., 2021) with 2,365 cM built from a backcross (BC_1_) population with 216 individuals and 8,402 SNPs retained with >80% call rate. Linkage group recombination sizes of our map were also similar to those reported by Yang et al. in the BC_1_ map (Spearman rank correlation 0.72), including the only large recombination gap in chromosome 15, starting at around the 40 cM position (Figure 1). This recombination gap was hypothesized to be due to reduced heterozygosity resulting from a selection sweep in favor of the allele transmitted from the Dwarf parent observed in the YANG genome assembly (Yang et al., 2021).

Although our map was built from a smaller number of individuals than the BC_1_ map used to anchor the YANG genome (Yang et al., 2021), those were derived from two recombined gametes from the F_1_ parents, therefore sampling twice as many meiotic events as in a BC1 population. The number of recombination events per meiosis captured in the mapping population is one of the main drivers of mapping accuracy, with an F_2_ being the most efficient configuration (Liu, 1998; Ferreira et al., 2006). Additionally, for improved linkage map quality, we decided to use only SNPs with call rate >95% aware of the potential reproducibility challenges of marker data derived from genotyping by sequencing methods in heterozygous genomes (Myles et al., 2010). From the raw data, 10,576 DArTSeq markers were retained at >95% call rate, and the main detractor of the final SNPs number was the test for the 1:2:1 segregation. This substantiates the variability in the F_1_ parents contributed by the Tall grandparental pollen pool. This observation indicates that a self-pollinated progeny from a single F_1_ plant would possibly allow mapping a considerably larger number of DArTSeq SNP markers.

### 4.2 The F_2_ linkage map provides an independent assessment of the properties of the coconut genome assemblies

The DArTseq method using the rare cutter SbfI combined to the frequent MseI enzyme proved effective for complexity reduction and enrichment of the sequenced pool for low copy genomic regions across most of the large coconut genome, generating a large number of segregating markers (Sansaloni et al., 2011). Our mapping and data quality control approach proved effective to order SNPs markers along the linkage groups into robust maps, highly colinear with the genome sequences (Table 1). The physical distance covered by our F_2_ linkage map was 1035.4 Mb on the YANG genome, slightly higher than the 1,020 Mb of the Yang et al. BC_1_ map, while on the WANG genome, the mapped SNPs covered 2,382 Mb, reaching a 99.27% coverage. As expected, very few segregating DArTSeq sequences and the corresponding SNPs were mapped to long physical stretches covering several hundred Mb of DNA of the reference-grade WANG genome assembly produced by nanopore long read sequencing (Figure 1). These regions were not sampled in the YANG genome, assembled exclusively with short read sequencing. The extensive genome gaps with no mapped SNPs and low gene density (Figure 2), likely correspond to heterochromatin, typically found at centromeres and telomeres, known to be relatively gene poor, mostly consisting of repetitive DNA sequences (Allshire and Madhani, 2018), where the DArTSeq method by principle tends not to sample restriction fragments. Performing linkage mapping in those heterochromatic regions will prove challenging, requiring approaches that can target the occasional low-copy regions scattered in the repetitive DNA. Using a selfed F_1_ progeny might prove valuable in this respect to sample much larger number of segregating DArTSeq SNPs, possibly targeting low-copy interspersed segments in these heterochromatic regions of the coconut genome.

The alignment of the linkage maps to the genome assembly indicated consistent ordering between the Joinmap estimated positions of the SNPs, and their relative physical order along the assembled chromosomes, with only sporadic inconsistencies seen most notably at the tip of some chromosomes (Figure 1). These localized inconsistencies could be due either to erroneous linkage map ordering or misassembled genome scaffolds. Historically, the majority of the inconsistencies between the physical and genetic map order pointed to errors in the physical map order (DeWan et al., 2002), making linkage maps effective tools to correct genome assemblies (Fierst, 2015). However, sequencing technologies and assembly algorithms have improved tremendously in recent years. The alignment of the linkage map to two independent genome assemblies, one of them built with long reads, indicates that those few SNPs that show inconsistent positions in relation to both genome assemblies, for example at the tips of chromosomes 8, 9 and 12, most likely were inaccurately mapped.

The largely consistent genetic to physical position of the mapped SNPs led us to carry out further analysis of this relationship for each coconut chromosome by estimating local recombination rates in cM/Mb, given by the slope of the curve built with graphical Marey maps. Additionally, the gene density histogram distribution annotated in the YANG genome was merged to the Marey maps for a combined view (Figure 2). In all chromosomes, the Marey maps showed genomic stretches of constrained recombination, where the recombination distance does not follow the linear increase in physical distance and gene density drops. This observation was evidently more distinct in the WANG genome Marey maps that included the heterochromatic regions of the genome. These regions are typically considered to be the putative location of centromeres. Considering the WANG chromosome numbering for now, these were located more toward the extremes of the chromosome (chromosomes 3, 7, 8, 9, 10, 12, 13 and 16) that would correspond to acrocentric or even telocentric chromosomes. In the other chromosomes these regions are found more toward the middle (Figure 2). Centromeres are essential for faithful segregation of chromosomes both in mitosis and meiosis and are typically rich in repetitive sequences in many eukaryotes (Achrem et al., 2020). Equivalent observations to ours on the putative centromere positions was reported in the YANG genome (Yang et al., 2021), establishing their correlation to the presence of repetitive rich DNA regions that were challenging to assemble. A few localized negative recombination rate estimates were also observed when the LOESS curve under passed the red dotted line, for example on chromosomes 7, 8, 9 and 12 (Figure 2). These localized negative estimates coincide with the inconsistent genetic to physical map alignments as described above and are usually located at the tip of chromosomes. All our results taken together, on the collinearity between our linkage map and both physical genomes provide an independent experimental validation of the genome assemblies currently available for coconut and establish the correspondence of chromosome numbering used in the two reports (Figure 2).

### 4.3 Colocation of a major effect QTL with the FLOWERING LOCUS T (FT) provides evidence for its role in early flowering

Little is known about the underlying mechanisms of early flowering in dwarf coconut. Knowledge on the complex physiology and genetics of flowering initiation has come mainly from annual plants including the model species *Arabidopsis* as well as crop plants, which typically flower and set seeds within a single year (Komeda, 2004; Roux et al., 2006). Less is known about flowering time in perennial plants which typically have distinguishable transition from a juvenile to an adult phase, the developmental stage that determines the emergence of flower meristems. It is now well established, however, that the formation of flower structures is largely conserved across flowering plants, annual and perennial, with relatively minor variations in the role of key developmental genes (Khan et al., 2014). Amongst the several genes identified as involved in regulation of flowering time, the expression of FLOWERING LOCUS T (FT) and its homologues has been shown to accelerate the onset of flowering in a number of plant species (Putterill and Varkonyi-Gasic, 2016). During the course of our study, we came across a transcriptomics report in coconut showing that the *FLOWERING LOCUS T* (FT) gene generates different transcripts in tall compared to dwarf coconut, with the shorter splice variant of FT present exclusively in dwarf varieties but absent in most tall coconuts (Xia et al., 2020). The authors suggested that this FT splice variant could be the putative causal mechanism for flowering time differentiation between the dwarf and tall coconut types, although they recommended further investigation and validation to confirm this effect. By collocating a phenotype-derived major QTL, detected agnostically to any prior gene-centered hypothesis, our study provides independent validation of the FT gene as the underlying major functional variants controlling flowering time between the two coconut varieties.

### 4.4 The *FT* locus likely controls other precocity traits in Dwarf coconut

A number of traits were targeted in previous QTL mapping studies in coconut, such as yield, fruit components, early germination and cuticular wax (Herrán et al., 2000; Ritter et al., 2000; Lebrun et al., 2001; Baudouin et al., 2006; Riedel et al., 2009). Flowering time, however could not be approached because it did not segregate in intermediate F_1_ hybrid families and less so in progeny of crosses between Tall cultivars. Two studies to date have reported on the patterns of genetic variation in F_2_ populations of coconut hybrids, although without attempting QTL mapping. Ample segregation was reported for a number of vegetative and morphological flowering traits, including height growth and the presence of a bole, for which the presence of at least one major gene each, with incomplete dominance was suggested (Namboothiri et al., 2011; Perera et al., 2016). Curiously, despite its importance as a trait directly linked to production, time to flower was not assessed. Our study is therefore the first one to tackle flowering time in coconut by a QTL mapping approach facilitated by the strong segregation seen in the F_2_. In our experiment we could not, however, measure height growth with confidence and include this trait in the mapping work. Recently a GWAS was carried out in a collection of Tall and Dwarf cultivars identifying a major effect QTL for height growth (Wang et al., 2021). The leading SNP for the GWAS hit was mapped to WANG chromosome 12, co-located in the promoter region of the gibberellin (GA) biosynthetic enzymes GA 20-oxidases (GA20ox) further backed by expression analysis, validating previous reports (Boonkaew et al., 2018). These results strongly support the role of GA20ox in controlling height growth difference between Tall and Dwarf coconuts. Taken together with the detection of a major flowering time QTL on chromosome 4 of WANG genome (chromosome 3 of YANG) (Figs 5 and 6) and its colocation with the FT gene, these results also suggest that height growth and flowering time in coconut involve different major effect loci in line with the early proposal of two independent major loci (Perera et al., 2016), although it cannot be ruled out that the FT gene is also involved in height growth (see below).

Petiole length (PL) and the presence of a bole (BOLE) were also assessed in our study. Both traits are typically used as morphological markers to identify dwarf plants (Batugal et al., 2009). In fact, PL and BOLE showed significant direct (0.466) and inverse (−0.502) correlation with flowering time (DLFS) (Figure 4). Major effect QTLs were also detected for these traits in the same exact region on chromosome 3 suggesting that the *FT* gene might also be affecting these traits (Figure 5). The *FT/TFL1* gene family has been extensively studied in flowering plants. It constitutes a major target of evolution in nature, as a single essential gene that has played a central role in plant diversification and adaptation (Pin and Nilsson, 2012). FT homologs have been identified in many species, demonstrating general conservation of functions across gymnosperms and angiosperms. It encodes a major florigen, a key flowering hormone in controlling flowering time. Studies in a number of species have revealed diverse roles of the FT/TFL1 gene family in plant developmental processes other than flowering regulation, with a distinct effect on stem and leaf development, mainly in the inhibition of stem and leaf growth (Wickland and Hanzawa, 2015; Freytes et al., 2021). It is therefore plausible that *FT* is directly affecting PL and the formation of a bole structure in coconut as well.

## 5. Conclusions: implications for coconut advanced breeding lines

From the conventional coconut breeding perspective, offspring of F_1_ hybrids would in principle not be desirable for commercial production due to the wide variation in yield traits among F_2_ coconut trees. Nevertheless, the two previous studies of F_2_ populations have reported individuals displaying high husked nut weights, indicating little or no inbreeding depression, likely resulting from the inbred nature of the ancestral Dwarf grandparent. These results indicated good potential of extracting recombined lines from F_2_ populations with desirable characters (Fernando and Perera, 1997; Namboothiri et al., 2011). Still, no developments have been made to implement this breeding strategy in coconut possibly due to the long time necessary to develop and eventually propagate such recombined lines.

The results of our study open a fundamentally new perspective in terms of marker assisted selection (MAS) in coconut breeding. Knowledge of the position and the genotypes at the SNP markers linked to the flowering time QTL would easily allow selecting hybrid F_2_ plants homozygous for the early flowering haplotype. Homozygosity for the favorable alleles at SNPs 100226499_C>T-34, 100287206_T>A-55 and 100273632_T>G-15, inherited from the Dwarf parent BGDJ caused a reduction in the time to flower around 400 days in our experiment. Our data also showed a significant negative correlation between the necessary days to flowering (DLFS) and NBF (Figure 4). Early flowering plants displayed some of the highest numbers of fruit per bunch and a separate QTL was mapped in chromosome 1 which could be selected by a set of linked SNPs flanking the lead SNP 100237259_A>G-48 (Table 2). This small set of SNPs could be used for high throughput inexpensive MAS screening of F_2_ plants at ultra-early stages of development or even at the embryo stage. Juvenile tissue of these selected individuals could be induced to somatic embryogenesis (Pérez-Núñez et al., 2006) or, more successfully, to shoot culture (Wilms et al., 2021) for clonal propagation. This would capture the full genetic value of these plants allowing regenerating thousands of plants indefinitely, providing the coconut industry with a solution for their current need of homogeneous quality planting material.

## Data availability statement

All genotypic and phenotypic data are included in the Supplementary Material. Further inquiries can be directed to the corresponding author.

## Author contributions

DG, WBSA and CAPP contributed to conception and design of the study. CAPP performed the field experimental work. WBSA carried out data analysis with input and help from DG. DG wrote the manuscript and all co-authors subsequently contributed to it by editing the final version.

## Acknowledgments

We would like thank the entire staff of field technicians EMBRAPA Tabuleiros Costeiros who were involved in establishing and managing the several field trials, and collecting and organizing the data used in this study. We thank Dr. Orzenil B. Silva-Junior and Dr. Roberto Togawa for assistance with the BLAST analysis against the YANG and Wang et al. genomes respectively.

## Institutional Review Board Statement

Not applicable.

## Informed Consent Statement

Not applicable.

## Supplementary material

**File S1.** DArTSeq SNP genotypic data for the full 3714 linkage mapped SNPs in the F_2_ population

**File S2.** Genotypic data for the 2952 DArTSeq SNP markers linkage mapped and physically mapped to unique positions on the coconut genome assemblies

**File S3.** BLAST results of the DArTSeq SNP corresponding sequences, target SNP type and position, against the YANG and WANG genomes.

**File S4.** Local recombination rates estimated following the Marey maps construction and LOESS smoothing between the linkage map and the two assembled genomes

**File S5.** Raw phenotypic data and estimated BLUPs following the mixed model analysis for the evaluated traits in the F_2_ population

**File S6.** QTL mapping results obtained with the composite interval mapping method

**File S7.** BLAST alignment results of the *Arabidopsis thaliana* putative flowering signals mediating protein FT (At1g65480) mRNA, complete cds against the YANG chromosome 3

**File S8.** BLAST alignment results of the *Arabidopsis thaliana* putative flowering signals mediating protein FT (At1g65480) mRNA, complete cds against the WANG chromosome 4

