## Supplementary material for "High density linkage to physical mapping in a unique Tall x Dwarf Coconut (*Cocos nucifera* L.) outbred F_2_ uncovers a major QTL for flowering time colocalized with the *FLOWERING LOCUS T (FT)*": File S7

Query= AY065378.1 Arabidopsis thaliana putative flowering signals mediating
protein FT (At1g65480) mRNA, complete cds

Length=840

>Cocos nucifera cultivar Hainan Tall coconut chromosome 3, whole genome shotgun sequence
Sequence ID: CM017874.1 Length: 82374452
Range 1: 70554038 to 70554249

Score:108 bits(119), Expect:3e-23,
Identities:151/212(71%), Gaps:0/212(0%), Strand: Plus/Plus

```
Query  392       GAGATTGTGTGTTACGAAAATCCAAGTCCCACTGCAGGAATTCATCGTGTCGTGTTTATA
451
                 ||||||||||| || || | |||| | ||  |    || || || ||  || | ||| | 
Sbjct  70554038  GAGATTGTGTGCTATGAGAGTCCACGGCCGGCGCTTGGCATCCACCGGTTCATCTTTGTG
70554097

Query  452       TTGTTTCGACAGCTTGGCAGGCAAACAGTGTATGCACCAGGGTGGCGCCAGAACTTCAAC
511
                  |||| |  ||||||||  |||| ||||||||||| || ||||||||||| || ||| ||
Sbjct  70554098  CTGTTCCAGCAGCTTGGGCGGCAGACAGTGTATGCCCCTGGGTGGCGCCAAAATTTCGAC
70554157

Query  512       ACTCGCGAGTTTGCTGAGATCTACAATCTCGGCCTTCCCGTGGCCGCAGTTTTCTACAAT
571
                 || || || ||||| ||  ||||||| |||||    || || || ||||| |  |  || 
Sbjct  70554158  ACCCGGGACTTTGCAGAACTCTACAACCTCGGATCACCAGTCGCAGCAGTCTATTTTAAC
70554217

Query  572       TGTCAGAGGGAGAGTGGCTGCGGAGGAAGAAG  603
                 || ||||| |||   |||| ||| || |||||
Sbjct  70554218  TGCCAGAGAGAGTCGGGCTCCGGCGGGAGAAG  70554249
```

Range 2: 73265231 to 73265450

Score:88.7 bits(97), Expect:3e-17,
Identities:155/222(70%), Gaps:3/222(1%), Strand: Plus/Plus

```
Query  392       GAGATTGTGTGTTACGAAAATCCAAGTCCCACTGCAGGAATTCATCGTGTCGTGTTTATA
451
                 ||||||||  |||| ||||  || |||||     |||| || || ||  | ||||||   
Sbjct  73265231  GAGATTGTAGGTTATGAAAGCCCTAGTCCGGTGTCAGGGATCCACCGCATGGTGTTTGCG
73265290

Query  452       TTGTTTCGACAGCTTGGCAGGCAAACAGTGTATGCACCAGGGTGGCGCCAGAACTTCAAC
511
                  |||| | |||| | |||||  |||  |||| |||| |||| |  ||| |  |||||||||
Sbjct  73265291  CTGTTCCAACAGTTAGGCAGAGAAAGCGTGTTTGCCCCAGAGATGCGGCCCAACTTCAAC
73265350
```

```
Query  512       ACTCGCGAGTTTGC-TGAGATCTACAATCTCGGCCTTCCCGTGGCCGCAGTTTTCTACAA
570
                 ||  |  | |||||  | || | || |||| || |  || || ||||| || | || |||
Sbjct  73265351  ACCAGGAATTTTGCACGGGAAC-ACTATCTGGGGCCACCGGTTGCCGCTGTCTACTTCAA
73265409

Query  571       TTGTCAGAGGGAGAGTGGCTGCGGAGGAAGAAGACTTTAGAT  612
                 ||| ||||||||   ||||| ||| || |||||| || ||||
Sbjct  73265410  TTGCCAGAGGGAATCTGGCTCCGGCGGTAGAAGA-TTCAGAT  73265450
```
