## Supplementary material for "High density linkage to physical mapping in a unique Tall x Dwarf Coconut (*Cocos nucifera* L.) outbred F_2_ uncovers a major QTL for flowering time colocalized with the *FLOWERING LOCUS T (FT)*": File S8

Query= AY065378.1 Arabidopsis thaliana putative flowering signals mediating
protein FT (At1g65480) mRNA, complete cds

>GWHBEBT00000004   OriSeqID=Chr04   Len=180429735
Length=180429735

 Score = 108 bits (119),  Expect = 7e-21
 Identities = 151/212 (71%), Gaps = 0/212 (0%)
 Strand=Plus/Minus

```
Query  392        GAGATTGTGTGTTACGAAAATCCAAGTCCCACTGCAGGAATTCATCGTGTCGTGTTTATA  
451
                  ||||||||||| || || | |||| | || |    || || || ||  || | ||| | 
Sbjct  164688301  GAGATTGTGTGCTATGAGAGTCCACGGCCGGCGCTTGGCATCCACCGGTTCATCTTTGTG  
164688242

Query  452        TTGTTTCGACAGCTTGGCAGGCAAACAGTGTATGCACCAGGGTGGCGCCAGAACTTCAAC  
511
                   |||| |  ||||||||  |||| ||||||||||| || ||||||||||| || ||| ||
Sbjct  164688241  
CTGTTCCAGCAGCTTGGGCGGCAGACAGTGTATGCCCCTGGGTGGCGCCAAAATTTCGAC  164688182

Query  512        ACTCGCGAGTTTGCTGAGATCTACAATCTCGGCCTTCCCGTGGCCGCAGTTTTCTACAAT  
571
                  || || || ||||| ||  ||||||| |||||    || || || ||||| |  |  || 
Sbjct  164688181  ACCCGGGACTTTGCAGAACTCTACAACCTCGGATCACCAGTCGCAGCAGTCTATTTTAAC  
164688122

Query  572        TGTCAGAGGGAGAGTGGCTGCGGAGGAAGAAG  603
                  || ||||| |||   |||| ||| || |||||
Sbjct  164688121  TGCCAGAGAGAGTCGGGCTCCGGCGGGAGAAG  164688090
```

 Score = 88.7 bits (97),  Expect = 7e-15
 Identities = 155/222 (70%), Gaps = 3/222 (1%)
 Strand=Plus/Plus

```
Query  392        GAGATTGTGTGTTACGAAAATCCAAGTCCCACTGCAGGAATTCATCGTGTCGTGTTTATA  
451
                  ||||||||  |||| ||||  || |||||     |||| || || ||  | ||||||   
Sbjct  168065258  
GAGATTGTAGGTTATGAAAGCCCTAGTCCGGTGTCAGGGATCCACCGCATGGTGTTTGCG  168065317

Query  452        TTGTTTCGACAGCTTGGCAGGCAAACAGTGTATGCACCAGGGTGGCGCCAGAACTTCAAC  
511
                   |||| | |||| | |||||  |||  |||| ||| |||| |  ||| |  |||||||||
Sbjct  168065318  
CTGTTCCAACAGTTAGGCAGAGAAAGCGTGTTTGCCCCAGAGATGCGGCCCAACTTCAAC  168065377

Query  512        ACTCGCGAGTTTGC-TGAGATCTACAATCTCGGCCTTCCCGTGGCCGCAGTTTTCTACAA  
570
                  ||  |  | |||||  | || | || |||| || |  || || ||||| || | || |||
```

Sbjct  168065378  ACCAGGAATTTTGCACGGGAAC-ACTATCTGGGGCCACCGGTTGCCGCTGTCTACTTCAA 168065436

Query  571        TTGTCAGAGGGAGAGTGGCTGCGGAGGAAGAAGACTTTAGAT  612
|  |  |  |  |  |  |  |  |  |  |  |  |  |  |  |  |  |  |  |  |  |  |  |  |  |  |  |  |  |  |  |  |
Sbjct  168065437  TTGCCAGAGGGAATCTGGCTCCGGCGGTAGAAGA-TTCAGAT  168065477
